# Wide scope analysis of bioactive lipids, including steroids, bile acids, and polyunsaturated fatty acid metabolites, in human plasma by LC/MS/MS

**DOI:** 10.1101/2023.04.13.536679

**Authors:** Kohta Nakatani, Yoshihiro Izumi, Hironobu Umakoshi, Maki Yokomoto-Umakoshi, Tomoko Nakaji, Hiroki Kaneko, Hiroshi Nakao, Yoshihiro Ogawa, Kazutaka Ikeda, Takeshi Bamba

## Abstract

Quantitative information on blood metabolites has the potential to be utilized in medical strategies such as early disease detection and prevention. Monitoring of bioactive lipids, such as steroids, bile acids, and polyunsaturated fatty acid (PUFA) metabolites, could be a valuable indicator for health status. However, a method for simultaneous measurement of these bioactive lipids has not been reported at present. Here, we report a liquid chromatography tandem mass spectrometry (LC/MS/MS) method that can simultaneously measure more than 140 bioactive lipids, including steroids, bile acids, and PUFA metabolites, from human plasma, and a sample preparation method for these targets. Protein removal in methanol precipitation and purification operations of bioactive lipids by solid-phase extraction improved the recovery of targeted compounds in human plasma samples, demonstrating the importance of sample preparation methods in a wide range of bioactive lipid analyses. Using the developed method, we measured plasma from healthy human volunteers and confirmed the presence of bioactive lipid molecules associated with sex differences and circadian rhythms. The practical bioactive lipid analysis method developed is expected to be applied to health monitoring and disease biomarker discovery for precision medicine.

## Introduction

If the health status can be objectively evaluated by the information of blood metabolites, it has the potential to be applied to medical strategies such as early detection and prevention of diseases (1). Metabolomics, which mainly focuses on comprehensive monitoring of hydrophilic metabolites, has been greatly applied to biomarker discovery (2), and the number of reports is increasing (3). However, despite the increase in biomarker research, the number of clinical applications is stagnant (3). Ongoing cohort studies have reported that concentrations of some amino acids increase with age and BMI (4), while other studies have reported no significant differences in amino acid levels with age or BMI and that smoking decreases amino acid levels (5). In these studies, although there are differences in sample size and race, it is assumed that there are many cross-reactions in hydrophilic primary metabolites due to various factors such as age, BMI, smoking status. As a result, the success of clinical applications may be very limited (6).

Steroids are a group of molecules that are biosynthesized from cholesterol mainly in the adrenal glands, transmitted via the blood to various organs, and exert their physiological activity by binding to membrane receptors and nuclear receptors (7). The measurement results of aldosterone [primary aldosteronism (8, 9)], cortisol [Cushing’s syndrome (10, 11), adrenal cortical carcinoma (12)], and progesterone [ovulation status (13)] are widely used in clinical practice. Bile acids are biosynthesized from cholesterol primarily in the liver, concentrated in the gallbladder, and excreted in the intestine. Excreted primary bile acids are either absorbed or circulate in the enterohepatic circulation after being converted to secondary bile acids by intestinal bacteria (14). In recent years, bile acids have attracted great attention because they have been reported to stimulate various immune systems (15).

Polyunsaturated fatty acid (PUFA) metabolites, such as eicosanoids and docosanoids, are biosynthesized from fatty acids cleaved from phospholipids that make up cell membranes by PLA2 (16, 17). It is also known to regulate inflammation and anti-inflammatory effects as immune cell receptors (16). Therefore, information on the in vivo profile of these bioactive lipids (i.e., steroids, bile acids, and PUFA metabolites) can be a promising indicator of health status.

When analyzing targeted endogenous metabolites from biological samples, it is important to apply appropriate sample preparation and analytical methods (18). As for analytical methods, Liquid chromatography mass spectrometry (LC/MS) is often used as an analytical method to analyze multiple compounds simultaneously, as in metabolomics (19, 20). However, it is still reportedly difficult to analyze a wide range of biological compounds from hydrophilic (e.g. ATP; log*P*_*ow*_, −5.5) to highly hydrophobic (e.g. TG 18:0/18:0/18:0; log*P*_*ow*_, 17.41) compounds with a single LC/MS technique. Therefore, a comprehensive analysis of a wide range of metabolome is underway by combining multiple measurement techniques and sample preparation methods (21).

Until now, LC/MS has been performed to measure metabolites in each category for research purposes, such as steroids (22), bile acids (23), and PUFA metabolites (24). However, no method has been reported to measure these compound groups across the board. Here, the log*P*_*ow*_ values for cortisol, cholic acid, and arachidonic acid, representative examples of steroids, bile acids, and PUFA metabolites, respectively, are 1.61, 2.02, and 6.98. Thus, from the perspective of the metabolome as a whole, these three groups are relatively close in physicochemical properties (25). Therefore, a detailed evaluation of the analytical system using a set of authentic standards may allow for the development of a simultaneous assay for these bioactive lipids.

On the other hand, the importance of sample preparation methods has recently been reevaluated, particularly by the Lipidomics Standard Initiative (LSI) and the International Lipidomics Society (iLS) (26). The role of sample preparation is to purify the target components and remove impurities. Failure to understand the role of sample preparation can lead to unexpected problems. For example, without proper pretreatment, impurities such as denatured proteins or precipitated highly hydrophobic lipids can accumulate on LC columns. Therefore, as the number of analyses increases, the pump pressure increases, which may reduce the reproducibility and accuracy of the analysis (27). Additionally, unwanted hydrophilic compounds can cause ion suppression of targeted compounds and reduce the sensitivity of LC/MS analysis, making proper sample preparation an essential step in obtaining high-quality data (28).

The purpose of this study is to propose a liquid chromatography tandem mass spectrometry (LC/MS/MS) method and sample preparation for comprehensive, simultaneous, and quantitative analysis of bioactive lipids, such as steroids, bile acids, and PUFA metabolites, for monitoring human health status. First, we optimized the multiple reaction monitoring (MRM) conditions for 145 bioactive lipids and stable-isotope-labeled internal standards (ISs) in triple-quadrupole mass spectrometry. Next, LC conditions for bioactive lipids were determined by examining the LC column and mobile phase conditions. The optimized LC/MS/MS method allows for the simultaneous analysis of 145 bioactive lipids. The sample preparation method was then evaluated based on the removal of impurities from plasma and the recovery rate of the targeted compounds. Optimized sample preparation and LC/MS/MS methods were used to measure bioactive lipids in human plasma of healthy volunteers and analyze sex differences and intra-day variations. The newly developed analytical method for bioactive lipids is the first report of a cross-sectional quantitative measurement of a group of compounds of steroids, bile acids, and PUFA metabolites.

## Materials and Methods

### Chemicals and reagents

LC–MS-grade water, acetonitrile, methanol, and 2-propanol were purchased from Kanto Chemical Co., Inc. (Tokyo, Japan). LC–MS-grade ammonium acetate was purchased from Merck (Darmstadt, Germany). LC–MS-grade acetic acid was purchased from Fujifilm Wako Pure Chemical Co. (Osaka, Japan). Authentic standards were obtained from Nacalai Tesque, Inc. (Kyoto, Japan), Fujifilm Wako Pure Chemical Co., and Merck. Authentic standards were obtained from Cayman Chemical Co. (Ann Arbor, MI), Cambridge Isotope Laboratories, Inc. (Tewksbury, MA), Steraloids Inc. (Newport, RI), Avanti Polar Lipids Inc. (Alabaster, AL), and Merck.

### Calculation of the structural properties

SMILES (Simplified Molecular Input Line Entry System) notation of all compounds extracted from PubChem database (25) was uploaded onto the online chemical database (OCHEM) web platform (29) for the calculation of log*P*_*ow*_, pKa, and pKb. All log*P*_*ow*_ were obtained with alvaDesc model, and pKa and pKb with the ChemaxonDescriptors model (Table S-1).

### Human plasma preparation

Plasma was collected from 8 healthy volunteers, 5 males and 3 females (Table S-2). Blood collection from males was performed in the morning (8:00–9:00), noon (12:00–13:00), and evening (17:00–18:00). Blood collection from females was performed in the morning (8:00–9:00). All plasma samples were collected before meals and centrifuged. The resulting supernatants were stored in a −80°C freezer until analysis. In the evaluation of the pretreatment method, equal amounts of mixed plasma samples from healthy subjects were used as quality control samples.

### Bioactive lipids extraction from plasma

The plasma was thawed at 4°C for approximately 12 h. A mixture of 50 μl of plasma, 500 μl of methanol, and 10 μl of IS containing 28 stable-isotope-labeled compounds in methanol (Table S-3) was vortexed for 1 min and then subjected to sonication for 5 min. The sample was incubated on ice for 5 min, followed by centrifugation at 4°C, 16,000 *g* for 5 min to precipitate proteins. The supernatant (500 μl) was transferred to a 2 ml Eppendorf tube and mixed with 1500 μl of water/formic acid (FA) (100/0.1, vol/vol) to obtain a sample for solid phase extraction (SPE) after protein removal by methanol precipitation. Another sample for SPE without protein removal by methanol precipitation was prepared by mixing 50 μl of plasma with 950 μl of water/methanol (75/25, vol/vol) and 10 μl of ISs in methanol, followed by vortexing for 1 min and then subjected to sonication for 5 min.

For SPE, an OASIS HLB 1 cc (30 mg) cartridge (Waters, Milford, MA) was first set in a vacuum manifold, and the cartridge was equilibrated by passing 1 ml of methanol/FA (100/0.1, vol/vol) and 1 mL of water/FA (100/0.1, vol/vol) serially. Then, the pre-prepared SPE samples were loaded onto the cartridge. Different combinations of solvents, such as water/FA (100/0.1, vol/vol), ethanol/water/FA (15/85/0.1, vol/vol/vol), methanol/water/FA (10/90/0.1, vol/vol/vol), methanol/water/FA (20/80/0.1, vol/vol/vol), methanol/water/FA (40/60/0.1, vol/vol/vol), and hexane, were examined as washing solvents. The detailed procedure was described in Table S-4. In the investigation of protein removal, the SPE cartridge was washed with water/FA (100/0.1, vol/vol), ethanol/water/FA (15/85/0.1, vol/vol/vol), and hexane in that order [described as methanol precipitation followed by SPE (MeP-SPE1) in Table S-4] was used. The optimized sample preparation procedure was MeP-SPE1, followed by recovering target bioactive lipids with 1 ml of methanol. The 1 ml of methanol in 1.5 ml Eppendorf tube was dried using a centrifugal evaporator and then reconstituted by vortexing with 60 μl of water/methanol (1/1, vol/vol). After being incubated on ice for 5 min, the samples were centrifuged at 4°C, 16,000 *g* for 5 min, and the 50 μl of resulting supernatant was transferred to a 0.3 ml Low Adsorption Vial QsertVial (Merck).

### Analytical conditions for LC/MS/MS

LC/MS/MS analyses were performed using a Nexera X2 UHPLC system (Shimadzu Co., Kyoto, Japan) coupled with an LCMS-8060 triple-quadrupole mass spectrometer (TQMS, Shimadzu Co.) with a heated electrospray ionization source. The LC mobile phase conditions used to evaluate the separation behavior were as follows: mobile phase (A), 5 mM ammonium acetate or 0.1% (vol/vol) acetic acid in water/acetonitrile (75/25, vol/vol); and mobile phase (B), 5 mM ammonium acetate or 0.1% (vol/vol) acetic acid in 2-propanol. The four LC columns used to compare bioactive lipids separation were as follows: Inertsil ODS-4; InertSustain C18; Inertsil ODS-HL; and Inertsil ODS-P [each, 2.1 mm inner diameter (i.d.) × 150 mm, 3 μm particle size, (GL Sciences Inc., Tokyo, Japan)]. The simplified gradient conditions used to compare separation behavior were as follows: 1% B, 0 min; 1– 99% B, 0–20 min; 99% B, 20–35 min; 99–1% B, 35–35.1 min; and 1% B, 35.1–45 min. The other LC conditions used for condition screening were as follows: injection volume, 10 μl; flow rate, 0.3 ml min^−1^; and column temperature, 40°C.

The optimized final LC analysis conditions used for plasma analyses were as follows: column, InertSustain C18; injection volume, 10 μl; flow rate, 0.3 ml min^−1^; column temperature, 50°C; mobile phase (A), 5 mM ammonium acetate in water/acetonitrile 75/25 (vol/vol); and mobile phase (B), 5 mM ammonium acetate in 2-propanol. The optimized gradient conditions were as follows: 1% B, 0 min; 1–38% B, 0–17 min; 38–99% B, 17–25 min; 99% B, 25–35 min; 99–1% B, 35–36 min; and 1% B, 36–46 min.

The MS conditions used for all analyses were as follows: nebulizer gas flow, 2 l min^−1^; heating gas flow, 10 l min^−1^; drying gas flow, 10 l min^−1^; heat block temperature, 400°C; desolvation line temperature, 250°C; and spray voltage, 4.0 kV for positive ion mode and −3.0 kV for negative ion mode. The MRM mode was applied to all targeted LC/MS/MS analyses. The MRM parameters were as follows: dwell time, 2 ms; pause time, 2 ms; and polarity switching time, 5 ms. The other MRM conditions, including the Q1 pre-bias, collision energy, and Q3 pre-bias of each metabolite, were optimized by flow injection analysis with standard solutions (1−100 μM) using LabSolution ver. 5.91 (Shimadzu Co.). The details of the optimized MRM parameters for the 145 targeted metabolites and 28 IS compounds were shown in Table S-3. To create calibration curves for each metabolite, standard solutions were prepared at concentrations of 0, 0.1, 1, 4, 10, 40, 100, 400, 1,000, 4,000, 10,000 nM. The LC/MRM data analysis was performed using LabSolutions, ver. 5.91 (Shimadzu Co.). The scan mode, with scan range *m/z* 100–1,000 for positive and negative respectively, was applied to estimate untargeted matrix constituents. The analytical platform for bioactive lipid analysis was controlled using LabSolutions (version 5.99 SP2, Shimadzu Co.)

### Data analysis

Data analysis for LC/MRM was performed using Multi-ChromatoAnalysT (Beforce Co., Fukuoka, Japan). Principle component analysis (PCA) was performed using MetaboAnalyst 5.0 (30). Spearman’s correlation analysis and other data analyses were performed in Excel 2016.

## Results and Discussion

### Calculation of Physicochemical Properties of Target Compounds and Optimization of MRM Conditions

In this study, we tried to construct a simultaneous and quantitative analytical method for a total of 145 unconjugated and conjugated forms of bioactive lipids (steroids, bile acids, and PUFA metabolites), including glucuronide, sulfate, glycine, and taurine conjugates (Table S-3).

The physicochemical properties such as hydrophobicity and charge characteristics are the major factors that affect the separation behavior in chromatography or the extraction and recovery efficiency from biological samples. By considering the physicochemical properties of the target metabolites, it is possible to propose a reasonable sample preparation and analytical method. Therefore, we obtained log*P*_*ow*_, pKa, and pKb of targeted metabolites using OCHEM, web platform for the calculation of various chemical information (29) (Table S-1). The range of log*P*_*ow*_, pKa, and pKb values for the targeted steroids, bile acids, and PUFA metabolites were 0.58-6.70, −1.75-19.78, and −7.48-9.31, respectively. These values were used for the consideration and optimization of the analytical and sample preparation methods.

LC/MS/MS in MRM mode has attracted attention for widely targeted metabolome analysis owing to its selectivity, high sensitivity, and good quantitative performance (19, 20). The MRM transitions (precursor ion, collision energy, product ion, and prequadrupole focusing voltages) of 145 metabolites and 28 stable isotope labeled compounds were optimized by flow injection analysis of each authentic standard, with up to two MRM transitions for each metabolite (Table S-3).

### Optimization of LC conditions

Reversed-phase liquid chromatography (RPLC) is suitable for separating hydrophobic compounds. In the present study, we defined the characteristics of four ODS columns with different hydrophobicity and stereoselectivity using the Tanaka method of silica ODS column characterization (31) (Table S-5). The gradient mode from water-acetonitrile mixture to 2-propanol, which has excellent elution power in RPLC, was used. However, the high viscosity of 2-propanol resulted in an increased column back pressure. Under the screening condition, while Inertsil ODS-4, InertSustain C18, and Inertsil ODS-HL showed no pressure issues (with maximum pressures of 42, 37, and 45 MPa, respectively), Inertsil ODS-P exceeded the pressure limit (>50 MPa). Therefore, Inertsil ODS-P was excluded from further consideration.

Following the analysis of the target compounds under six different LC conditions (using three columns and two set of mobile phases, ammoniuim acetate and acetic acid additives), a total of 128 compounds were detected as single peaks. The peak information, such as retention time (RT) and peak width, for each detected compound was summarized in Table S-6. To evaluate the effect of six different LC analytical conditions with three different columns and two additives on the retention behavior of bioactive lipids, PCA analysis was performed using RTs for each compound (Figure 1 and Table S-7). The scores plot showed that each LC condition could be plotted using PC1 (67.9%) and PC2 (21.4%), and the differences in RTs were clearly reflected in the conditions. Among the ODS columns studied, the type of additive had a significant impact. Of the 66 compounds with a loading 1 value of less than 0, only 14 compounds (21.2%) had a pKa less than 5, whereas of the 62 compounds with a loading 1 value of more than 0, 58 compounds (93.5%) had a pKa less than 5. It was considered that the difference in additives between ammonium acetate and acetic acid would greatly affect the retention of acidic compounds. Because the acetic acid additive condition produced broad tailing peaks for many compounds, we selected ammonium acetate as the additive in this study. It was highlighted that additives are important for the chromatographic performance, especially for compounds with a wide range of physicochemical properties, such as steroids, bile acids, and PUFA metabolites.

**Figure 1.**
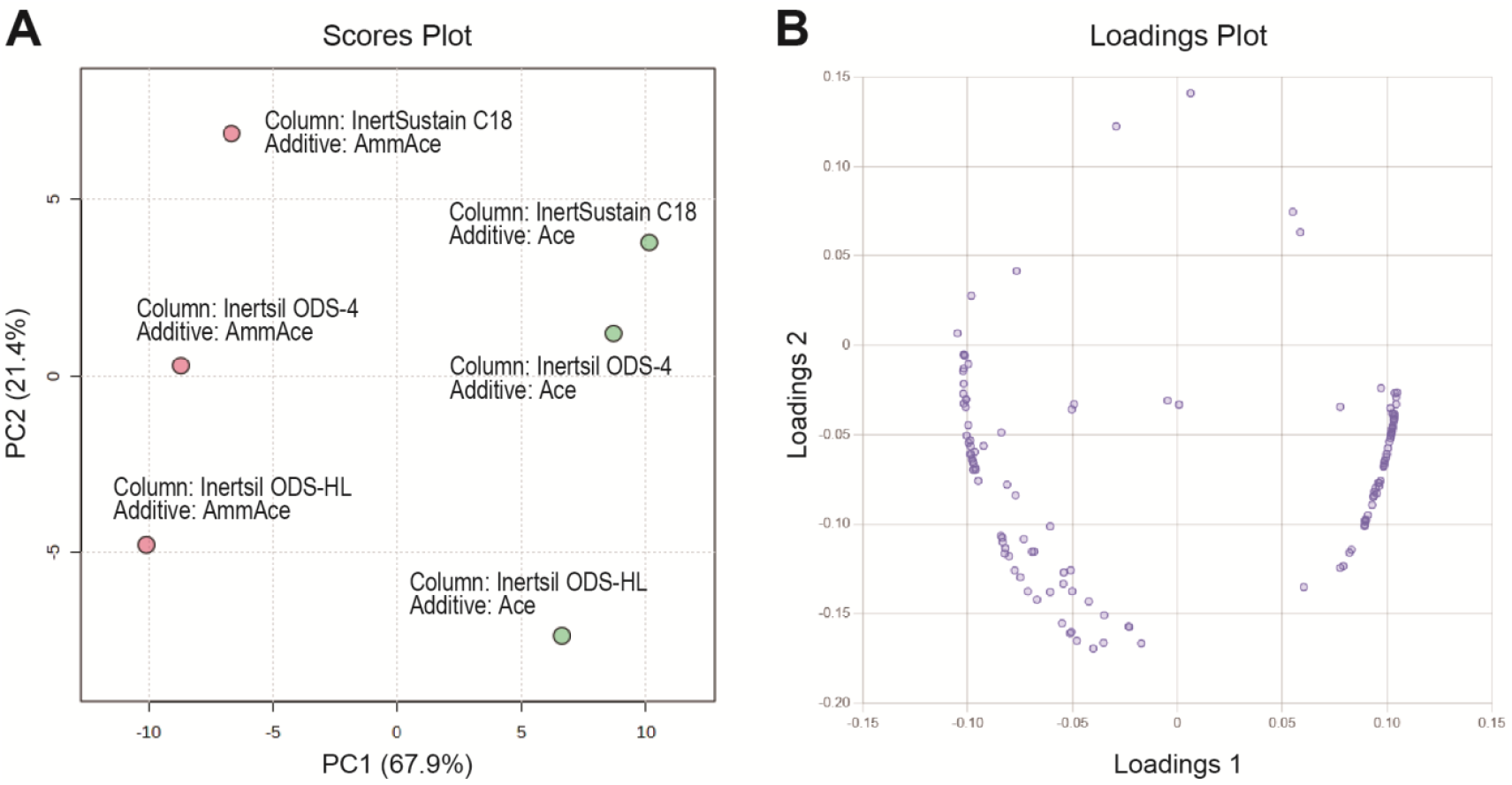
PCA results for characterization of LC conditions with bioactive lipids. The retention times of bioactive lipids detected consistently across all screening conditions were used. Scores plot is shown in **A**, where red dots represent ammonium acetate additive condition, and green dots represent acetic acid additive condition. Loadings plot is shown in **B**.

We conducted Spearman’s correlation analysis between the loading values and log*P*_*ow*_ of each compound. The correlation coefficient for loadings 1 was 0.21, while that for loadings 2 was 0.47, indicating a correlation between loading 2 and log*P*_*ow*_. Therefore, it was considered that PC2 could be partially explained by hydrophobicity. It is reasonable that each LC condition was plotted in order on the PC2 axis based on the types of ODS columns. However, no differences in the separation performance of the targeted isomers were observed among Inertsil ODS-4, InertSustain C18, and Inertsil ODS-HL. Therefore, we selected InertSustain C18, which allowed for stable measurements with the lowest column back pressure.

Finally, we optimized the LC conditions to improve chromatographic separation and enhance the stability of the analysis. Specifically, in order to improve the separation of weakly retained compounds, the gradient slope in the early half of the analysis was adjusted to be more gradual, and the column temperature was raised from 40°C to 50°C to reduce column backpressure. The chromatogram obtained under the optimized conditions was shown in Figure 2. The results of the analysis validation were described in Table S-3. The developed analytical method enabled the measurement of wide-range of steroids, bile acids, and PUFA metabolites.

**Figure 2.**
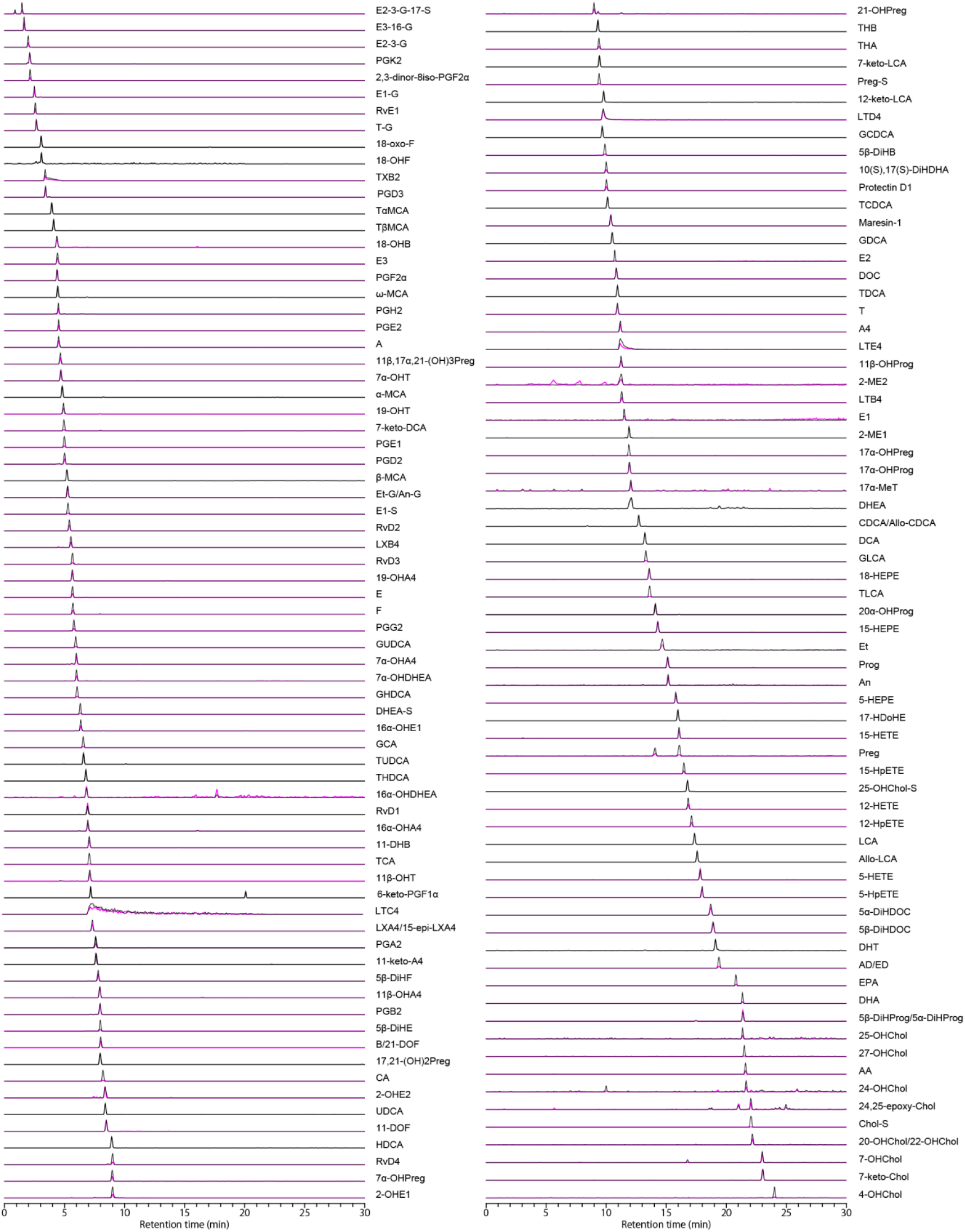
Separation overview of optimized LC/MS/MS method. The chromatograms in black represent quantitative MRM transitions, while the chromatograms in pink represent qualitative MRM transitions. All compound abbreviations are defined in Table S-3.

### Evaluation of sample preparation method for wide-targeted bioactive lipids analysis

The evaluation of sample preparation methods is crucial for achieving stable and continuous measurements of biological samples, especially when conducting simultaneous analysis of a large number of compounds. The role of sample preparation method is to remove impurities and extract/recover target components. In this study, we assumed impurities to be proteins, hydrophilic metabolites, and highly hydrophobic metabolites such as phospholipids and neutral lipids. First, we investigated the removal of impurities.

Removal of proteins: proteins can be removed as precipitates by mixing them with organic solvents, trapping them in SPE cartridges, or washing them away with solvents in SPE process. We evaluated protein removal using three methods: methanol precipitation (MeP), SPE, and MeP followed by SPE (MeP-SPE1) (Figure S-1, Table S-4). The SPE method used in this study was based on the previously reported method for extracting eicosanoids (32). As a result, all three methods (MeP, SPE, and MeP-SPE1) showed removal efficiencies of 97% or higher for proteins (97.7%, 98.5%, and 99.6%, respectively) (Figure 3A). These methods were effective enough to remove proteins sufficiently.

**Figure 3.**
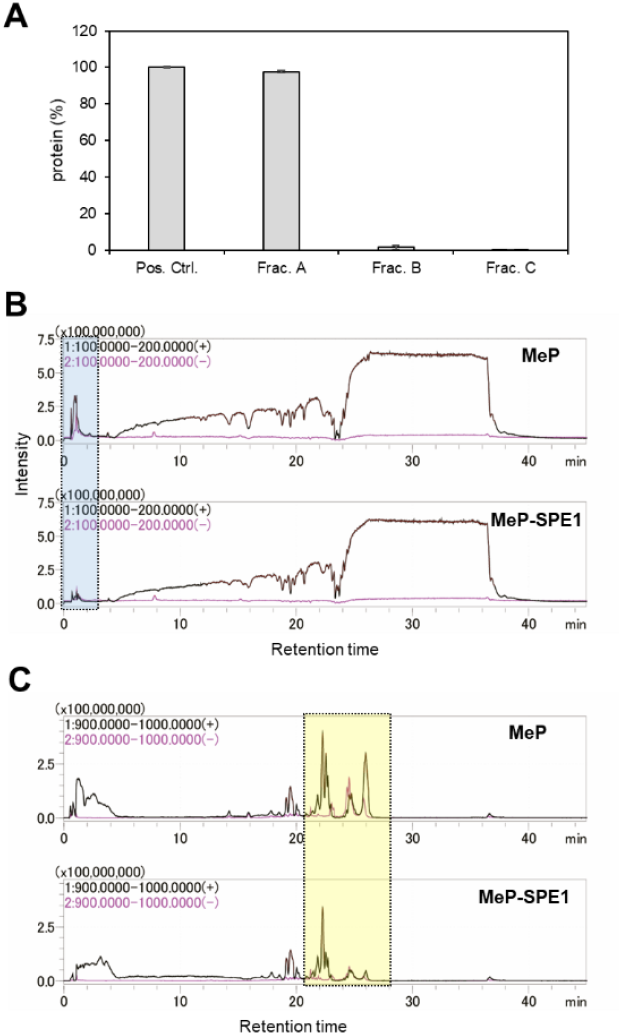
Evaluation of impurity removal for proteins, hydrophilic compounds, and hydrophobic compounds. **A**. Evaluation of protein removal. Frac. A showed the precipitate from methanol suspension of plasma, Frac. B showed the elution fraction from SPE, and Frac. C showed the elution fraction using SPE after methanol suspension. As a positive control, plasma was directly subjected to BCA assay. **B**. Evaluation of the removal of low molecular weight hydrophilic compounds. MS chromatograms of human plasma in scan mode of *m/z* 100-200 with different sample preparation methods was described. In RPLC, hydrophilic compounds elute in the early retention time range, as highlighted in blue. **C**. Evaluation of the removal of highly hydrophobic compounds. MS chromatograms of human plasma in scan mode of *m/z* 900-1000 with different sample preparation methods was described. In RPLC, highly hydrophobic compounds elute in the later retention time range, as highlighted in yellow.

Removal of hydrophilic compounds: hydrophilic compounds such as standard amino acids with monoisotopic mass less than 186.1, are considered to be removed during the washing step with an aqueous solvent in SPE. In RPLC/MS using an ODS column, hydrophilic compounds are typically not retained and are eluted immediately after sample injection. Therefore, to confirm the removal of hydrophilic compounds by SPE, the SPE procedure was performed by changing the set of washing solvents (see details in Table S-4). The MeP-treated supernatant and the eluted fraction after MeP-SPE were measured using the ODS-RPLC/MS scan mode (Figure S-2). By comparing the MeP-treated supernatant and MeP-SPE elution fraction, a decrease in the ion intensity of total ion current (TIC) in the range of *m/z* 100-200 and retention time of less than 2.5 was observed in the MeP-SPE elution fraction (Figure 3B). The results suggested that SPE effectively removed hydrophilic compounds. On the other hand, no significant difference in the removal of hydrophilic compounds was observed among the six SPE procedures studied (Figure S3-4). The full scan mode of the quadrupole MS was considered to have limitations in capturing detailed changes.

Removal of highly hydrophobic compounds: highly hydrophobic compounds, including phospholipids (e.g., PI 20:4_22:6, monoisotopic mass 930.5) and neutral lipids (e.g., TG 56:6, monoisotopic mass 906.8), are thought to be removed by the washing process of SPE or by being trapped in the SPE cartridge. By comparing the MeP-treated supernatant and MeP-SPE elution fraction, a decrease in ion intensity of TIC in the range of *m/z* 900-1000 with a retention time of 20 minutes or more was observed in the MeP-SPE elution fraction (Figure 3C), suggesting the removal of highly hydrophobic compounds by SPE. The six SPE procedures investigated did not show significant differences in the removal of highly hydrophobic compounds, probably for the similar reason to hydrophilic compounds (Figure S3-4). To summarize the removal of impurities, it was possible to remove most, or a certain amount, of impurities such as proteins, hydrophilic compounds, and highly hydrophobic compounds by MeP-SPE procedures.

Next, we investigated the recovery rates of target compounds. We prepared 28 stable isotope-labeled ISs of steroids, bile acids, and PUFA metabolites, and calculated the recovery rates of these compounds when subjected to MeP-SPE either alone or with plasma (Table 1). In the evaluation of ISs without plasma, MeP-SPE1 had the highest number of compounds (26) with ISs recoveries of more than 70%. Compounds that showed poor recovery rates overall were taurocholic acid-d5 (TCA-d5), taurolithocholic acid-d5 (TLCA-d5), and cholesterol sulfate-d7 (Chol-S-d7), which were taurine and sulfate conjugates with a sulfo group in their substructure and exhibited high acidity (pKa of −1.06, −0.84, and −1.36, respectively). Additionally, TCA-d5 and TLCA-d5 had high RSDs of over 30% in recovery rate. When evaluating the recovery of ISs without plasma, compounds with a sulfo group as a substructure had poor recovery rates and repeatability.

**Table 1.** Recovery rate of steroids, bile acids, and PUFA metabolites.

| Sample |  | IS |  |  |  |  |  |  |  |  |  |  |  |  |  | plasma with IS |  |  |  |  |  |  |  |  |  |  |  |
| --- | --- | --- | --- | --- | --- | --- | --- | --- | --- | --- | --- | --- | --- | --- | --- | --- | --- | --- | --- | --- | --- | --- | --- | --- | --- | --- | --- |
| Method |  | MeP |  | MeP-SPE1 |  | MeP-SPE2 |  | MeP-SPE3 |  | MeP-SPE4 |  | MeP-SPE5 |  | MeP-SPE6 |  | MeP-SPE1 |  | MeP-SPE2 |  | MeP-SPE3 |  | MeP-SPE4 |  | MeP-SPE5 |  | MeP-SPE6 |  |
| Compound | Group | Ave.<br>(%) | RSD<br>(%) | Ave.<br>(%) | RSD<br>(%) | Ave.<br>(%) | RSD<br>(%) | Ave.<br>(%) | RSD<br>(%) | Ave.<br>(%) | RSD<br>(%) | Ave.<br>(%) | RSD<br>(%) | Ave.<br>(%) | RSD<br>(%) | Ave.<br>(%) | RSD<br>(%) | Ave.<br>(%) | RSD<br>(%) | Ave.<br>(%) | RSD<br>(%) | Ave.<br>(%) | RSD<br>(%) | Ave.<br>(%) | RSD<br>(%) | Ave.<br>(%) | RSD<br>(%) |
| B-d8 | steroid | 100 | 2.3 | 81.3 | 3.1 | 77.3 | 3.9 | 83.2 | 4.8 | 84.3 | 2.2 | 86.2 | 2.8 | 85.8 | 4.7 | 87.9 | 2.3 | 79.0 | 11.3 | 78.0 | 0.8 | 56.2 | 15.1 | 80.9 | 5.2 | 94.1 | 0.5 |
| 18-OHB-d4 | steroid | 100 | 1.8 | 76.0 | 1.5 | 77.7 | 1.7 | 80.4 | 0.4 | 83.8 | 0.9 | 84.7 | 1.3 | 82.4 | 1.5 | 38.0 | 3.4 | 38.8 | 2.6 | 39.3 | 5.5 | 38.7 | 7.1 | 35.7 | 4.2 | 38.2 | 2.0 |
| DOC-d8 | steroid | 100 | 3.8 | 89.7 | 3.5 | 85.4 | 3.1 | 94.2 | 3.0 | 96.8 | 3.3 | 97.5 | 3.8 | 93.6 | 5.5 | 91.0 | 0.2 | 74.6 | 9.9 | 83.8 | 8.2 | 58.4 | 18.2 | 83.2 | 9.1 | 97.4 | 1.1 |
| 11-DOF-d5 | steroid | 100 | 5.0 | 87.0 | 3.3 | 85.2 | 1.3 | 89.9 | 8.5 | 93.4 | 4.9 | 90.6 | 0.2 | 89.0 | 3.2 | 82.0 | 3.4 | 70.4 | 9.1 | 74.4 | 2.1 | 54.0 | 14.1 | 76.7 | 5.2 | 85.0 | 0.9 |
| F-d4 | steroid | 100 | 3.3 | 72.1 | 4.0 | 71.7 | 2.6 | 75.5 | 1.8 | 83.6 | 3.1 | 83.2 | 7.1 | 85.2 | 3.7 | 43.6 | 4.9 | 35.8 | 3.0 | 35.0 | 3.6 | 28.7 | 4.9 | 34.0 | 6.1 | 44.6 | 8.6 |
| E-d8 | steroid | 100 | 5.1 | 85.4 | 4.4 | 83.6 | 1.4 | 89.9 | 3.7 | 88.9 | 1.6 | 87.9 | 1.3 | 87.1 | 3.4 | 57.9 | 10.0 | 49.2 | 5.3 | 45.3 | 3.0 | 35.1 | 7.5 | 45.2 | 3.1 | 53.1 | 3.0 |
| T-d3 | steroid | 100 | 4.4 | 89.4 | 1.0 | 77.3 | 2.4 | 90.9 | 6.1 | 88.2 | 1.8 | 95.5 | 4.7 | 92.0 | 5.3 | 96.6 | 4.1 | 79.5 | 12.1 | 89.7 | 5.6 | 60.1 | 20.1 | 86.8 | 4.8 | 105.0 | 5.1 |
| Chol-S-d7 | steroid | 100 | 24.4 | 18.6 | 11.4 | 8.8 | 9.3 | 4.0 | 3.9 | 1.8 | 0.7 | 3.5 | 3.6 | 9.6 | 12.4 | 24.2 | 4.2 | 45.3 | 13.0 | 31.5 | 25.8 | 26.4 | 10.3 | 45.7 | 10.9 | 10.6 | 6.1 |
| CA-d4 | bile acid | 100 | 6.8 | 78.2 | 1.4 | 76.6 | 4.3 | 76.8 | 4.0 | 71.9 | 0.6 | 76.5 | 2.8 | 77.5 | 3.0 | 72.7 | 3.0 | 67.6 | 11.1 | 64.8 | 0.5 | 40.2 | 13.7 | 55.3 | 8.3 | 55.4 | 0.4 |
| CDCA-d4 | bile acid | 100 | 4.0 | 99.8 | 4.3 | 95.3 | 3.8 | 93.4 | 6.0 | 87.7 | 6.1 | 89.2 | 2.8 | 92.0 | 4.9 | 85.7 | 3.0 | 75.5 | 11.4 | 72.5 | 2.7 | 43.9 | 16.2 | 66.6 | 6.2 | 65.5 | 2.7 |
| DCA-d4 | bile acid | 100 | 3.3 | 103.6 | 0.5 | 97.9 | 1.0 | 97.1 | 4.4 | 87.3 | 2.1 | 90.8 | 3.5 | 92.2 | 4.1 | 98.4 | 9.1 | 91.3 | 16.0 | 85.2 | 8.9 | 55.5 | 19.6 | 88.5 | 1.8 | 74.1 | 2.7 |
| LCA-d4 | bile acid | 100 | 9.9 | 141.1 | 9.8 | 140.6 | 15.0 | 127.7 | 9.4 | 109.8 | 8.8 | 119.9 | 0.9 | 120.1 | 11.2 | 124.2 | 14.1 | 136.6 | 32.4 | 97.9 | 37.5 | 68.7 | 27.4 | 107.0 | 8.6 | 66.3 | 7.3 |
| GCA-d4 | bile acid | 100 | 7.5 | 81.7 | 1.6 | 88.8 | 12.2 | 75.7 | 3.6 | 71.1 | 0.7 | 72.8 | 3.2 | 74.3 | 4.7 | 60.7 | 4.4 | 58.2 | 8.1 | 50.5 | 4.2 | 33.8 | 9.9 | 44.4 | 7.6 | 43.7 | 1.5 |
| GDCA-d4 | bile acid | 100 | 7.8 | 96.4 | 1.5 | 97.0 | 5.8 | 90.7 | 2.4 | 84.0 | 4.1 | 86.9 | 1.2 | 90.4 | 5.6 | 89.0 | 10.8 | 87.5 | 12.1 | 68.9 | 3.9 | 42.9 | 15.6 | 62.9 | 8.7 | 57.0 | 1.3 |
| GLCA-d4 | bile acid | 100 | 8.1 | 107.5 | 4.3 | 134.0 | 21.9 | 95.0 | 7.0 | 86.2 | 5.7 | 89.0 | 2.5 | 87.5 | 2.6 | 88.4 | 12.7 | 85.2 | 13.7 | 64.7 | 10.1 | 43.3 | 14.7 | 63.7 | 4.7 | 53.2 | 2.2 |
| TCA-d5 | bile acid | 100 | 11.4 | 86.7 | 47.4 | 62.0 | 46.4 | 35.4 | 33.6 | 11.9 | 4.7 | 33.6 | 46.2 | 54.5 | 42.4 | 31.9 | 3.9 | 35.0 | 7.4 | 16.2 | 9.3 | 6.1 | 5.7 | 21.0 | 6.4 | 19.8 | 13.7 |
| TLCA-d5 | bile acid | 100 | 4.7 | 75.7 | 53.3 | 53.7 | 47.0 | 16.8 | 21.6 | 2.2 | 1.1 | 19.9 | 32.5 | 34.4 | 47.9 | 26.9 | 6.1 | 27.0 | 5.4 | 14.5 | 0.8 | 8.1 | 2.2 | 15.8 | 0.7 | 11.4 | 1.4 |
| PGE2-d4 | PUFA | 100 | 5.3 | 72.1 | 2.2 | 76.4 | 4.8 | 69.3 | 3.6 | 68.1 | 1.4 | 74.0 | 1.3 | 74.3 | 7.0 | 61.9 | 7.1 | 50.5 | 6.0 | 49.9 | 5.3 | 28.6 | 11.7 | 43.7 | 12.8 | 40.4 | 3.4 |
| PGD2-d4 | PUFA | 100 | 5.1 | 70.9 | 2.2 | 77.4 | 7.1 | 72.1 | 1.5 | 70.4 | 1.9 | 71.5 | 1.1 | 76.2 | 5.2 | 122.0 | 18.1 | 145.5 | 26.0 | 125.5 | 12.0 | 69.4 | 28.3 | 119.1 | 0.7 | 92.8 | 5.4 |
| PGF2α-d4 | PUFA | 100 | 4.4 | 65.2 | 2.7 | 69.1 | 5.1 | 66.7 | 0.6 | 67.6 | 3.1 | 68.2 | 1.5 | 71.7 | 5.3 | 49.1 | 4.2 | 41.1 | 9.2 | 38.2 | 3.4 | 21.4 | 8.1 | 32.0 | 11.0 | 33.5 | 0.4 |
| TXB2-d4 | PUFA | 100 | 3.8 | 74.9 | 0.6 | 76.3 | 1.6 | 75.3 | 2.3 | 73.2 | 0.6 | 78.1 | 0.2 | 79.0 | 6.0 | 71.0 | 5.5 | 63.7 | 9.7 | 61.1 | 3.0 | 36.4 | 13.4 | 52.5 | 11.8 | 54.8 | 1.8 |
| LTD4-d5 | PUFA | 100 | 8.7 | 81.4 | 1.5 | 82.2 | 6.3 | 79.3 | 3.5 | 76.7 | 3.8 | 67.9 | 1.7 | 76.5 | 7.8 | 66.8 | 11.8 | 67.4 | 14.3 | 50.6 | 10.3 | 35.9 | 13.6 | 50.6 | 14.2 | 40.0 | 0.3 |
| LTB4-d4 | PUFA | 100 | 4.9 | 84.4 | 1.9 | 84.1 | 2.5 | 80.4 | 2.4 | 80.9 | 3.1 | 82.5 | 3.2 | 86.6 | 2.9 | 79.3 | 7.3 | 67.2 | 13.8 | 62.8 | 5.8 | 40.4 | 14.8 | 63.4 | 8.3 | 61.3 | 0.3 |
| 15-HETE-d8 | PUFA | 100 | 12.0 | 122.5 | 6.7 | 108.8 | 9.8 | 104.4 | 2.0 | 98.0 | 5.5 | 100.3 | 6.1 | 105.1 | 4.9 | 107.6 | 15.4 | 104.7 | 29.2 | 87.0 | 18.7 | 49.2 | 20.7 | 84.3 | 2.7 | 69.2 | 2.5 |
| 15-HEPE-d5 | PUFA | 100 | 6.4 | 124.1 | 9.7 | 114.2 | 13.9 | 115.2 | 6.8 | 106.2 | 2.6 | 108.2 | 2.0 | 110.8 | 4.4 | 119.0 | 12.9 | 98.4 | 23.6 | 95.8 | 8.1 | 49.8 | 18.7 | 87.0 | 1.4 | 84.7 | 2.7 |
| DHA-d5 | PUFA | 100 | 17.3 | 148.9 | 21.4 | 109.6 | 25.3 | 125.7 | 12.4 | 119.8 | 9.6 | 123.8 | 9.5 | 136.8 | 1.6 | 56.0 | 0.8 | 86.2 | 4.4 | 50.8 | 33.4 | 59.7 | 16.1 | 59.2 | 14.1 | 30.6 | 5.6 |
| AA-d8 | PUFA | 100 | 14.5 | 176.6 | 16.4 | 137.7 | 26.0 | 159.7 | 9.7 | 154.5 | 2.8 | 165.2 | 2.6 | 168.7 | 1.8 | 85.6 | 6.3 | 120.3 | 10.4 | 76.2 | 47.1 | 75.0 | 22.3 | 91.8 | 10.0 | 47.5 | 7.5 |
| EPA-d5 | PUFA | 100 | 9.8 | 149.0 | 11.9 | 121.2 | 25.4 | 126.0 | 11.7 | 124.6 | 6.1 | 133.0 | 3.6 | 140.3 | 7.3 | 76.0 | 5.2 | 107.4 | 9.5 | 63.5 | 37.8 | 71.5 | 22.5 | 87.6 | 7.8 | 49.0 | 6.9 |
| Recovery > 70% |  | 28 |  | 26 |  | 24 |  | 23 |  | 23 |  | 23 |  | 25 |  | 17 |  | 15 |  | 11 |  | 2 |  | 11 |  | 7 |  |
| RSD < 30% |  | 28 |  | 26 |  | 26 |  | 27 |  | 28 |  | 26 |  | 26 |  | 28 |  | 27 |  | 24 |  | 28 |  | 28 |  | 28 |  |

On the other hand, when ISs were added to plasma, the recovery rates tended to be lower than those obtained from the ISs without plasma. For example, the recovery rate of cortisol-d4 (F-d4) for MeP-SPE1 was 72.1% for ISs alone, but 43.6% for ISs with plasma. This was considered to be due to ion suppression caused by the plasma matrix. Moreover, when plasma and ISs were extracted together, there was a tendency for compounds with poor repeatability without plasma to show improved repeatability. For example, the repeatability of TCA-d5 and TLCA-d5 in MeP-SPE1 was 47.4% and 53.3%, respectively, with ISs alone, but improved to 3.9% and 6.1%, respectively, when extracted with plasma. The phenomenon of changes in elution behavior due to the loose binding of some matrix components to the analytes, previously suggested in chromatography (33), may partially explain the observed difference in repeatability between SPE with and without plasma matrix. The RSDs of ISs recovery with plasma matrix were less than 30% for all ISs in MeP-SPE1 (with a maximum RSD of 18.1% for PGD2-d4). Therefore, we decided to use MeP-SPE1 for plasma analysis in this study. Although further investigations on the extraction methods are needed to improve the recovery rate, the combination of developed LC/MS/MS and sample preparation method can provide quantitative values even for compounds with low recovery rates by using corresponding ISs for normalization.

At last, we state the accuracy levels of the quantitative values that could be obtained with our method. In LC/MS, using stable isotope dilution method allows for correcting factors affecting the quantification such as extraction efficiency and ionization efficiency, resulting in more accurate quantitation (34). However, when the number of targets to be measured exceeds 150, as is the case in this study, it is not practical to prepare stable isotope-labeled ISs for all targets due to limitations in running costs and commercial availability. The analytical method includes quantification values corrected using stable isotope-labeled ISs that correspond to target compounds as well as those with similar chemical properties but not necessarily identical to the target compounds. Improving the quantification of all measurement targets is a future challenge.

In summary, this study reevaluated the extraction methods with the expansion of measurement targets for steroids, bile acids, and PUFA metabolites. The removal of non-target proteins, hydrophilic compounds, and highly hydrophobic compounds and recovery rates of target compounds were examined. As a result, the LC/MS/MS method combined with MeP-SPE1 made it possible to measure a diverse set of bioactive lipids, including steroids, bile acids, and PUFA metabolites.

### Application to plasma bioactive lipid profiles in human healthy volunteer

Using the optimized method, we measured human plasma samples. The samples were obtained from eight healthy volunteers, including five males and three females. All blood was collected before meals, and female samples were collected in the morning, while male samples were collected at three time points: morning, noon, and evening. Detailed sample information is described in Table S-2.

As a result of measurement, we successfully measured a total of 46 bioactive lipids (19 steroids, 20 bile acids, and 7 PUFA metabolites) (Table S-8). We investigated the sex differences in the measured bioactive lipids using the morning samples. While metabolites that do not show sex differences like arachidonic acid (AA) were observed, the presence of metabolites that show sex differences, including sex hormones such as progesterone (Prog) for females and testosterone (T) for males, was suggested (Figure 4AB). This suggests the importance of exploring gender-specific metabolites beyond sex hormones. This suggests the importance of considering the existence of metabolites that differ by sex beyond the sex hormones, for example in biomarker discovery. Although Prog showed a large standard deviation, this is thought to be due to the influence of the menstrual cycle on female hormones.

**Figure 4.**
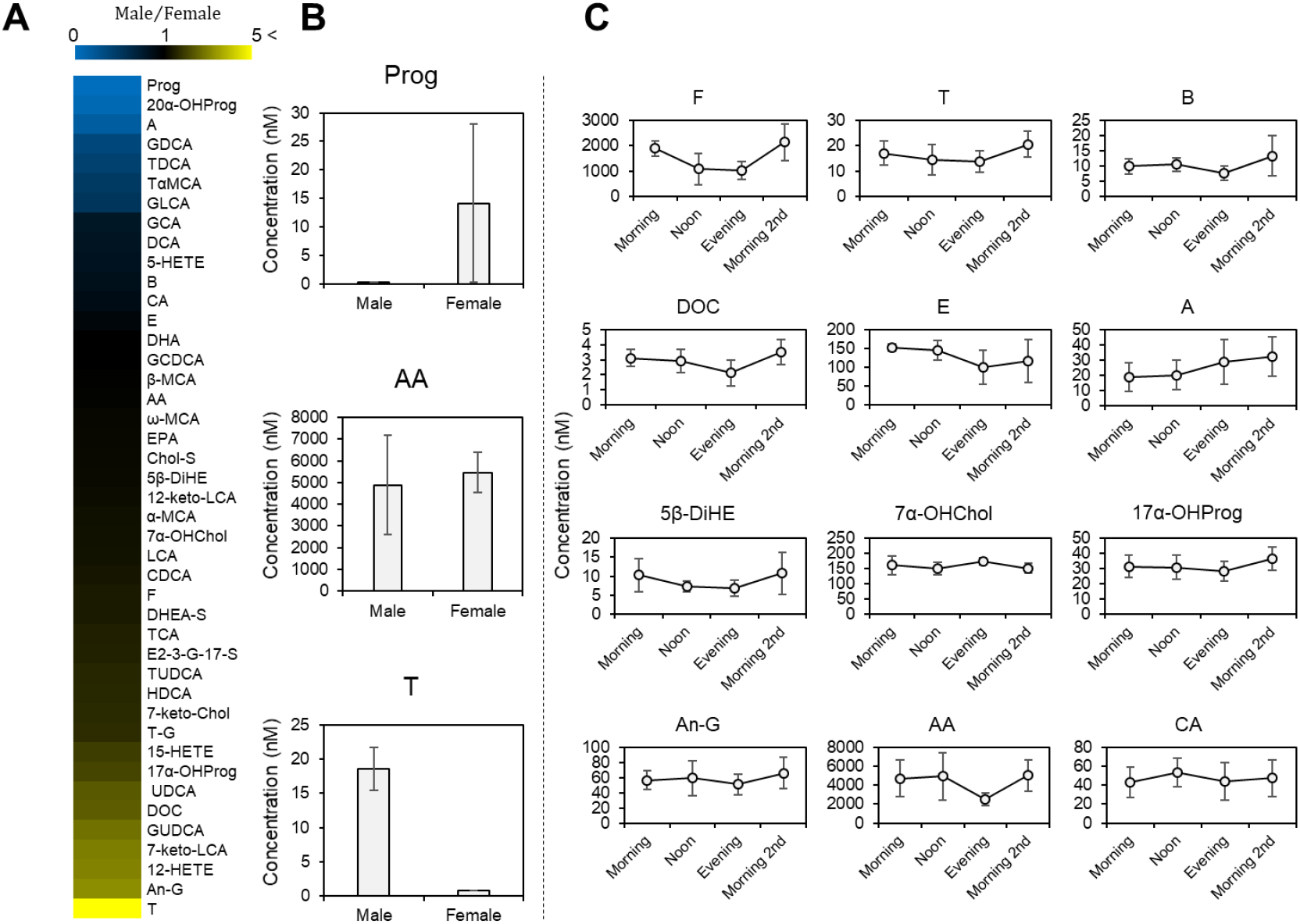
Investigation of sex differences and circadian rythms using plasma analysis from human healthy voluntters. **A**. Analysis of sex differences. The ratio of detected bioactive lipids in males (n=5) and females (n=3) was calculated. Those with a ratio of 1 are shown in black, those with a ratio of 0 (indicating more in females) are shown in blue, and those with a ratio of 5 or more (indicating more in males) are shown in yellow. **B**. Bar graph of representative bioactive lipids in the sex difference analysis. **C**. Analysis of circadian rythms. Male plasma was collected on a specific days in the morning (n=5), noon (n=5), and evening (n=4), and on the other days in the moring (n=7).

Next, we investigated the intra-day variation using male samples collected in the morning, noon, and evening. Cortisol (F) has been reported as a metabolite that exhibits a circadian rhythm, with high levels in the morning and low levels in the evening (35). In our result, it was possible to capture the circadian rhythm of F, which gradually decreased from morning to evening and returned to high levels the other morning (Figure 4C). Furthermore, similar intra-day variation was observed in other steroid hormones except for aldosterone (A) (Figure 4C). In addition, AA, which is grouped as PUFA metabolites, showed a tendency to have high levels in the morning and noon, while the cholic acid (CA), which is grouped as bile acids, showed high levels in the noon (Figure 4C). In this study, we revealed the presence of circadian rhythms in various bioactive lipids, in addition to the known circadian rhythm like F. This also suggests the importance of considering circadian rhythms in biomarker discovery. These findings are expected to provide useful information to increase the resolution of analysis in future biomarker discovery.

## Conclusion

We have successfully developed the LC/MS/MS method that enabled simultaneous measuring a wide range of bioactive lipids, including steroids, bile acids, and PUFA metabolites. We have also constructed the sample preparation method suitable for the simultaneous analysis of a large number of active lipids in human plasma by an effective combination of protein precipitation with methanol and purification by SPE. Using the developed sample preparation and analytical methods, we measured bioactive lipids in the plasma of human healthy volunteers, revealing the presence of sex differences and circadian rhythms in various bioactive lipids. The methods for measuring bioactive lipids that we have developed, along with the insights gained from measuring bioactive lipids in healthy plasma, are expected to contribute to improving the resolution of future biomarker discovery.

## Data Availability statement

Essential data are contained in the manuscript. Supporting data are provided as Supplemental Materials.

## Acknowledgments

This work was partly performed in Medical Research Center Initiative for High Depth Omics, and Cooperative Research Project Program of the Medical Institute of Bioregulation, Kyushu University.

## Footnotes

### Author Contribution

Y.I. and T.B. equally contributed to this work. K.N., Y.I., and T.B. designed the study. H.U., M. Y-U., H.K., H.N., Y.O. collected blood samples. K.N. and T.N. performed the experiments and data analysis. K.N., Y.I., and T.B. wrote the manuscript, and other authors contributed to manuscript editing.

---

**Figure S-1.**
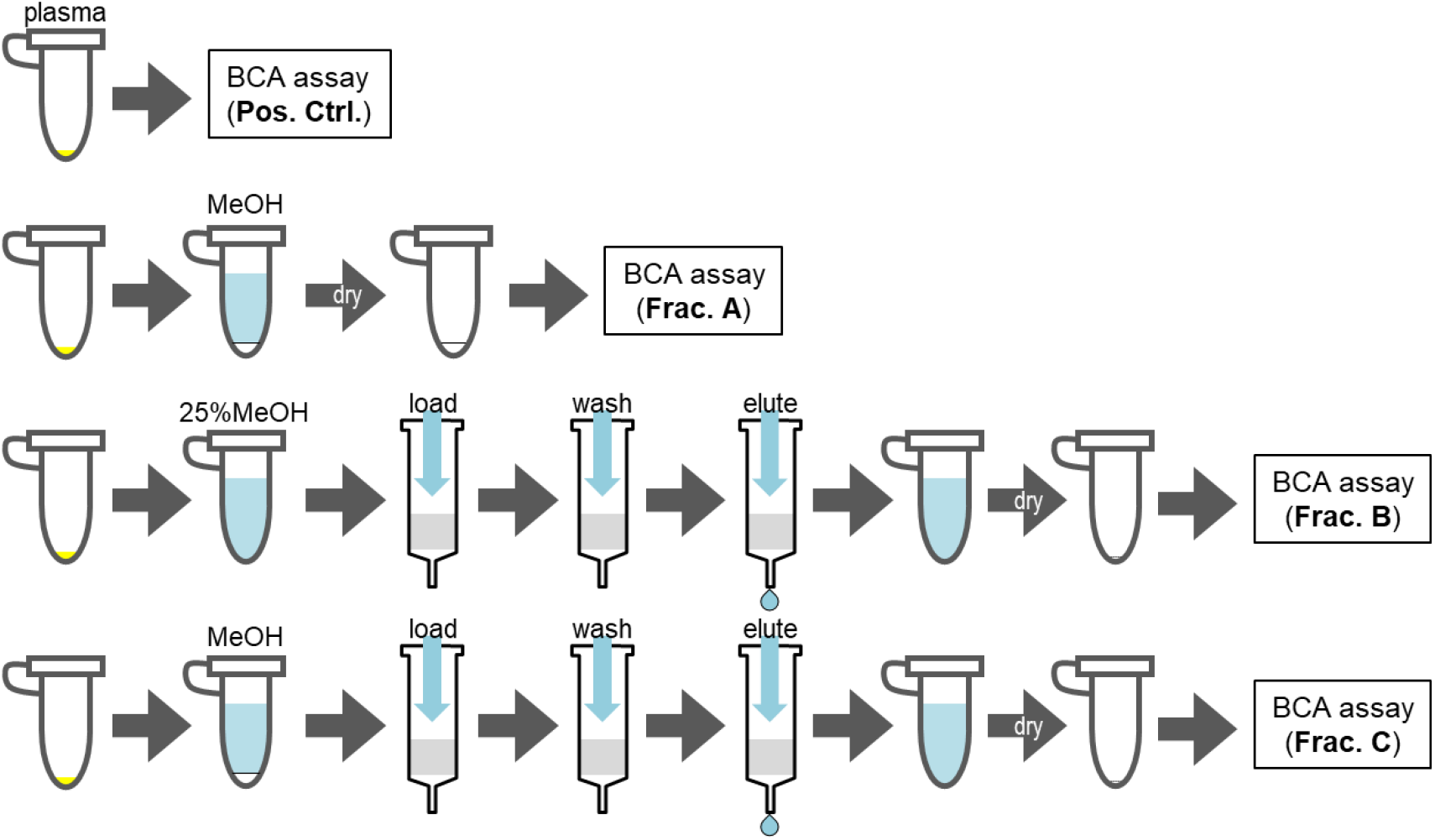
Overview of sample preparation procedure for evaluation of protein removal.

**Figure S-2.**
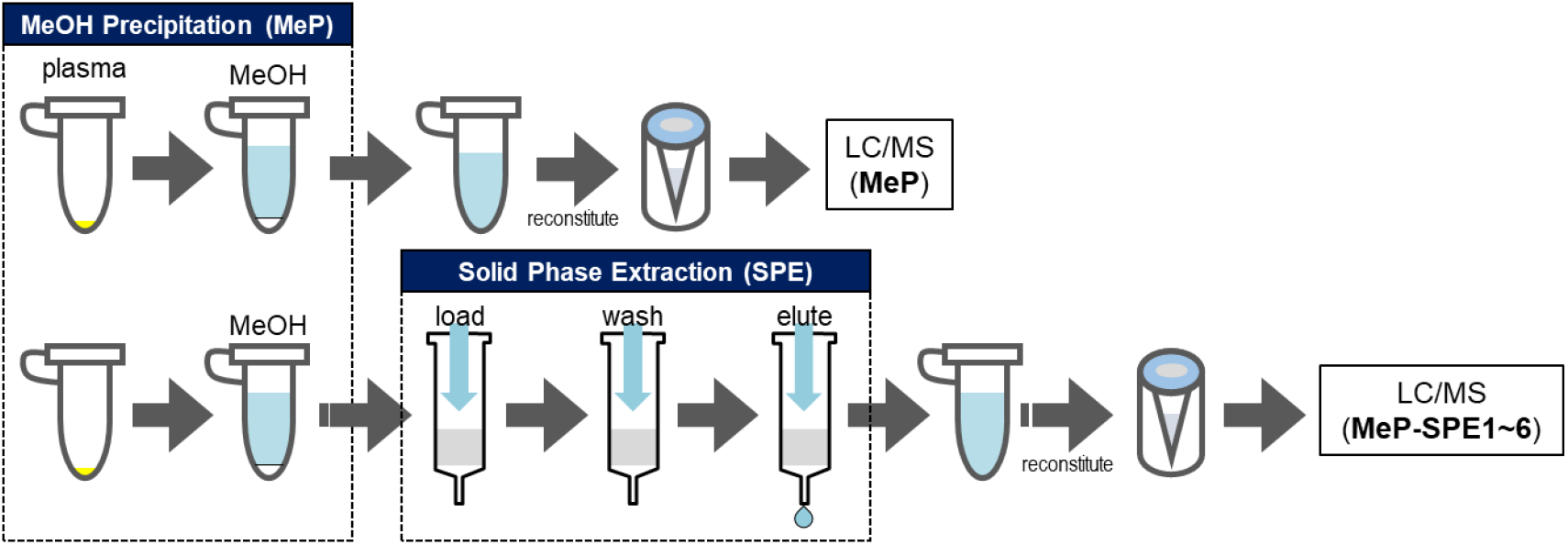
Overview of sample preparation procedure for evaluation of removal of impurities and recovery of targeted bioactive lipids.

**Figure S-3.**
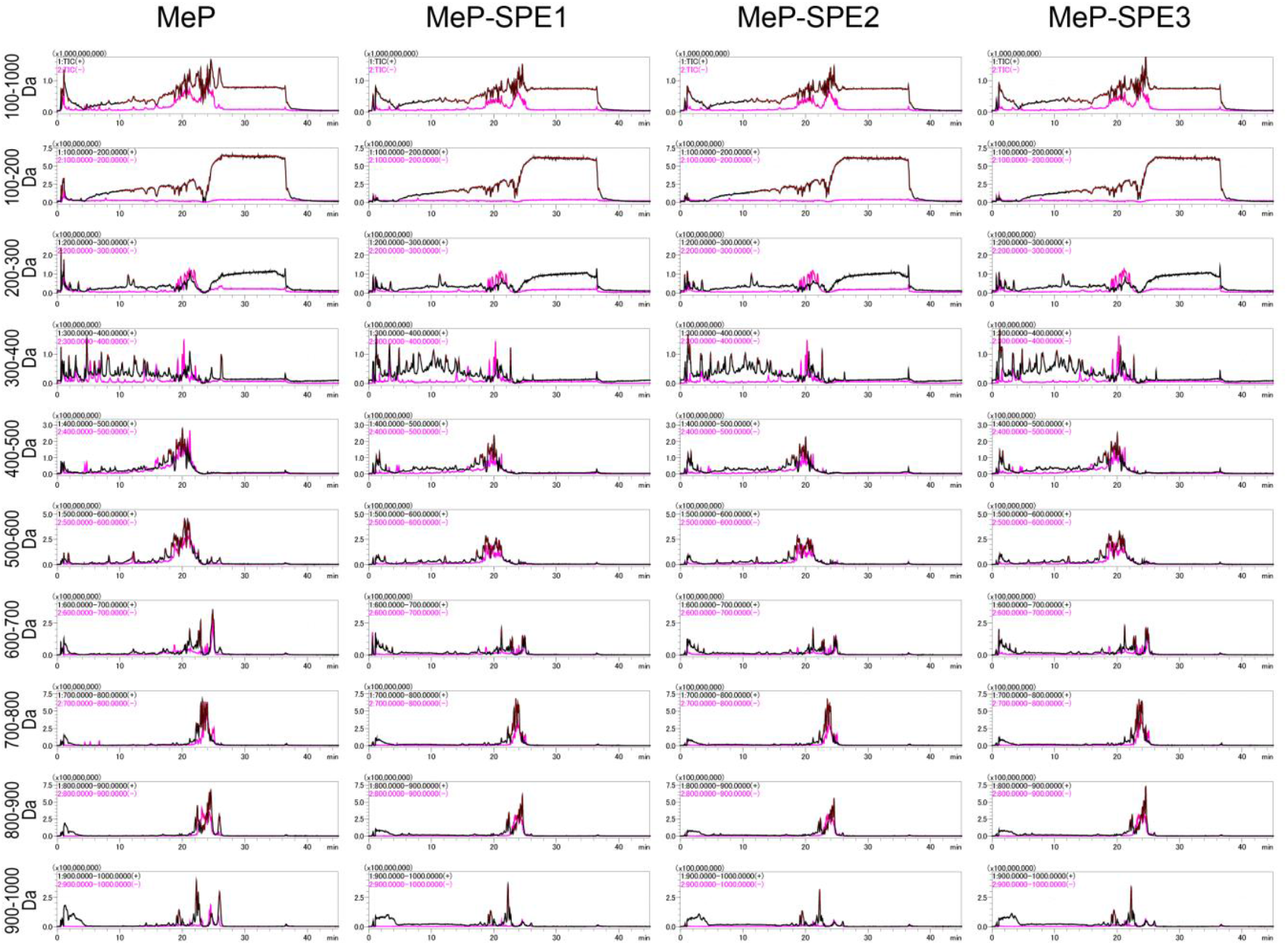
MS chromatograms of human plasma in scan mode with different sample preparation methods.

**Figure S-4.**
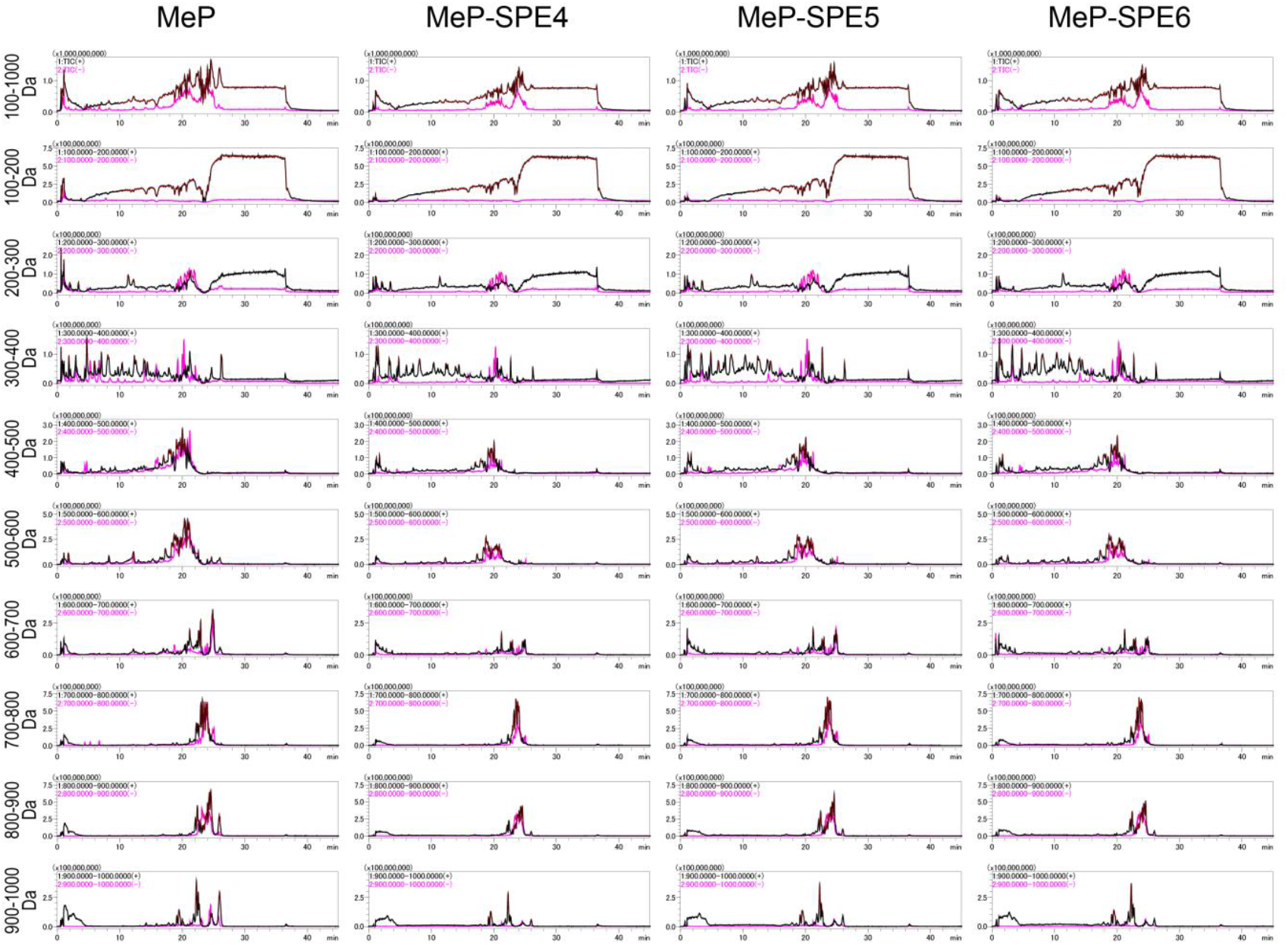
MS chromatograms of human plasma in scan mode with different sample preparation methods.

## Tables

Table S-1. Physicochemical properites of steroids, bile acids and PUFA metabolites in our library.

Table S-2. Sample information used in this study.

Table S-3. Database IDs, MRM conditions, and analytical validation of targeted LC/MS/MS method for steroids, bile acids and PUFA metabolites.

Table S-4. Solvents and order of use in solid phase extraction.

Table S-5. Information on the LC column investigated in this study.

Table S-6. Result of LC condition screening for steroids, bile acids, and PUFA metabolites.

Table S-7. Loadings of PCA analysis and representative chemical properties.

Table S-8. Results of bioactive lipid analysis in plasma of human healthy volunteers.

